# Host genetic associations with the gut microbiota in HIV-1-infected subjects: a pilot exploratory study

**DOI:** 10.1101/427922

**Authors:** Yolanda Guillén, Marc Noguera-Julian, Javier Rivera, Maria Casadellà, Muntsa Rocafort, Mariona Parera, Beatriz Mothe, Josep Coll, Jorge Saz, Jordi Navarro, Manuel Crespo, Eugènia Negredo, Christian Brander, Maria Luz Calle, Bonaventura Clotet, Roger Paredes

## Abstract

The impact of host genetics on gut microbial dynamics is debated. No study to date has investigated the possible role of host genetics in shaping the gut microbiota in HIV-1 infected subjects. With the aim of generating preliminary data to inform future host genetic studies, we performed an exploratory host exome analysis of 147 subjects either infected or at risk of becoming infected with HIV-1 from the MetaHIV cohort in Barcelona. Using a DNA microarray chip, we sought to identify host genetic variants associated to three specific microbial features with a potentially inheritable component, and which were previously found to be associated with gut dysbiosis in HIV infection, i.e.: gut enterotype, presence of methanogenic archaea and microbial gene richness. After correction for multiple comparisons, we did not observe any statistically significant association between the host’s genetic landscape and the explored gut microbiome traits. These findings will help design future, adequately-powered studies to assess the influence of host genetics in the microbiome of HIV-1-infected subjects.

## Introduction

The human gut microbiota performs essential functions to human health^1^, from nutrient digestion to immune homeostasis^2^. Perturbations of gut microbial configurations thus impact human health. They may occur due to a wide range of environmental factors, including diet^3,4^, diseases^5-7^, medication intake^8^, and are even influenced by age, geography^9^ and sexual behavior^10^. Over 20% of the inter-person microbiome variability is indeed associated with factors related to diet, drugs and anthropometric measurements^11^.

The host’s genetic background shapes the structure and influences the dynamics of the niche inhabited by the intestinal flora^12^. However, the influence of host genetics on the human microbiome is more debated^13^. To date, microbiome-wide association studies (MGWAs) integrating the effect of both environmental and host’s intrinsic factors on the microbiota have yielded disparate results ^14,15^. Most candidate genes associated with microbiota configurations participate in pathways involved in host immunity, microbial sensing, sugar digestion and host dietary preferences^16-18^. The abundance of intestinal *Bifidobacterium* has been associated with SNPs in the lactase-coding gene (LCT) in independent cohorts^13,19,20^. Moreover, an heritable component has been attributed to certain gut microbial taxa like Christensenellaceae^21^ and Methanobrevibacter^22^.

Understanding the reciprocal interaction between HIV-1 infection, the host immune system and the intestinal microbiota might provide clues to cure HIV-1 infection. In a crossectional study in 156 European subjects with different HIV-1 phenotypes (MetaHIV cohort) ^23,24^, we previously showed that men who have sex with men (MSM) were enriched in *Prevotella* and, after controlling for HIV-1 risk group, there was no consistent microbial dysbiosis pattern discernible by 16S rRNA sequencing^25^. Using shotgun metagenomics analyses in the same cohort, we identified a strong, dose-dependent relationship between gut microbial richness and nadir CD4+ T-cell counts^24^, which is the major surrogate marker of excess mortality, systemic inflammation and clinical complications in chronically HIV-1-infected subjects^26,27^. In our study, changes in microbial richness reflected multidimensional modifications in microbial species composition and functions, including increases in reactive oxygen and nitrogen-resistant species (ROS/RNS) like the Gram-negative generalists *Bacteroides* and Proteobacteria, and decreases in ROS/RNS-sensitive Gram-positive syntrophic microbes, including hydrogen-consuming methanogenic archaea.

A yet unanswered question is whether the host genetic background could have any influence in such gut microbiome shifts. To the best of our knowledge, no study has investigated the possible role of host genetics in the shaping the gut microbiota in HIV-1 infected subjects. With the aim of generating preliminary data to inform future larger studies, we performed a pilot, exploratory exome analysis of the MetaHIV cohort seeking to identify host genetic variants associated to 3 specific microbial features previously found to be important in this setting, and with a potential inheritable component, i.e.: gut enterotype^28^, presence of methanogenic archaea^22^ and microbial gene richness^29^.

## Results

### Population stratification in the MetaHIV cohort

We analyzed the exomes of 147 subjects from the MetaHIV cohort ^23,24^ using the Illumina Infinium CoreExome microarray, which covers 547,644 putative functional exonic variants (Table 1). Our study included 30 (20%) females and 117 (80%) males, most of them Caucasian (79.6%). The gut microbiota of these participants had been characterized using 16S rRNA gene amplicon sequencing^23^ and whole metagenome shotgun sequencing^24^ in previously published studies. Using 16S rRNA, 61 individuals from our cohort (41.5%) were classified as Bacteroides-rich, whereas 86 subjects (58.5%) were categorized as Prevotella-rich. Methanogenic archaea were detectable in 89 participants (60.5%).

**Table 1.**
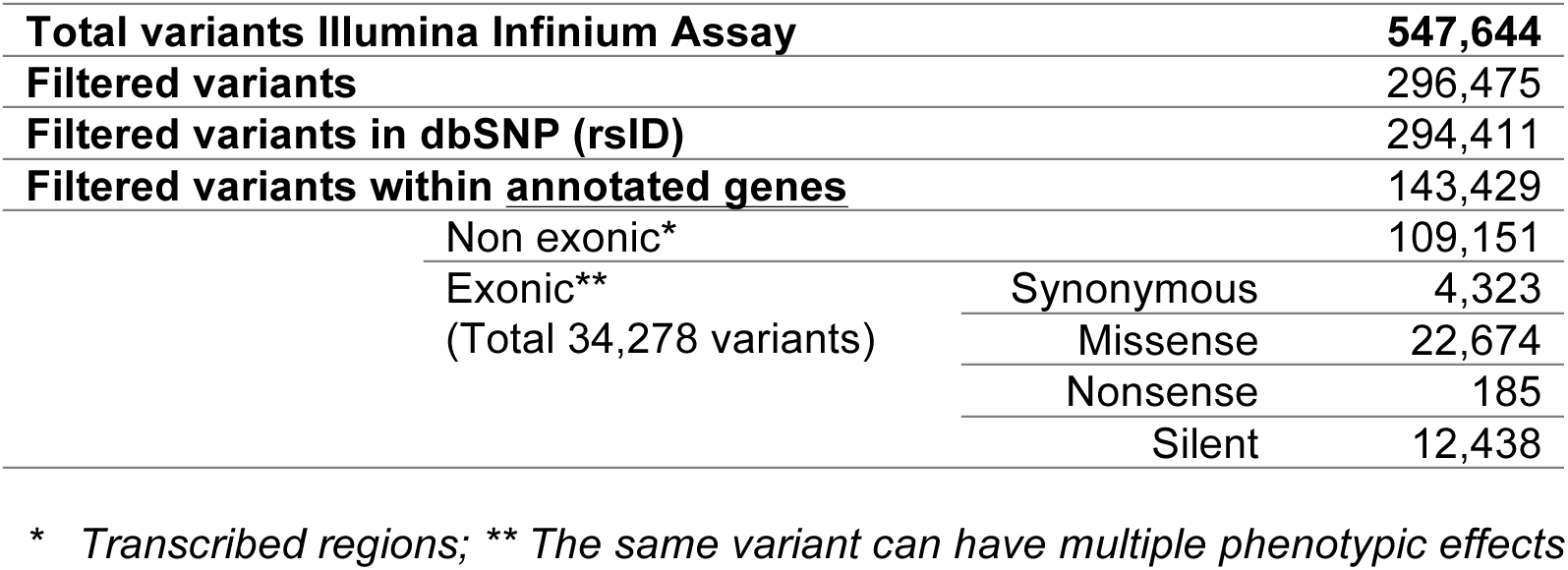
Human genetic variants analyzed with the Human Infinium^®^ CoreExome-24v1-0 BeadChip microarray and the Infinium^®^ HTS Assay Protocol Guide

From all variants covered by the Illumina^®^ microarray, 296,475 passed the pre-specified quality filters and were thus selected for further analyses. From these, 294,411 (99.3%) were found in dbSNP^30^, and 143,429 (48.4%) were located within an annotated human gene (Table 1). A multidimensional scaling (MDS) plot based on identity-by-state (IBS) distances showed that our multi-ethnic cohort was stratified by genetic ancestry (Figure 1a). As expected, females and males also showed a different pattern of missing variants as estimated by an identity-by-missingness (IBM) matrix (Figure 1b).

**Figure 1.**
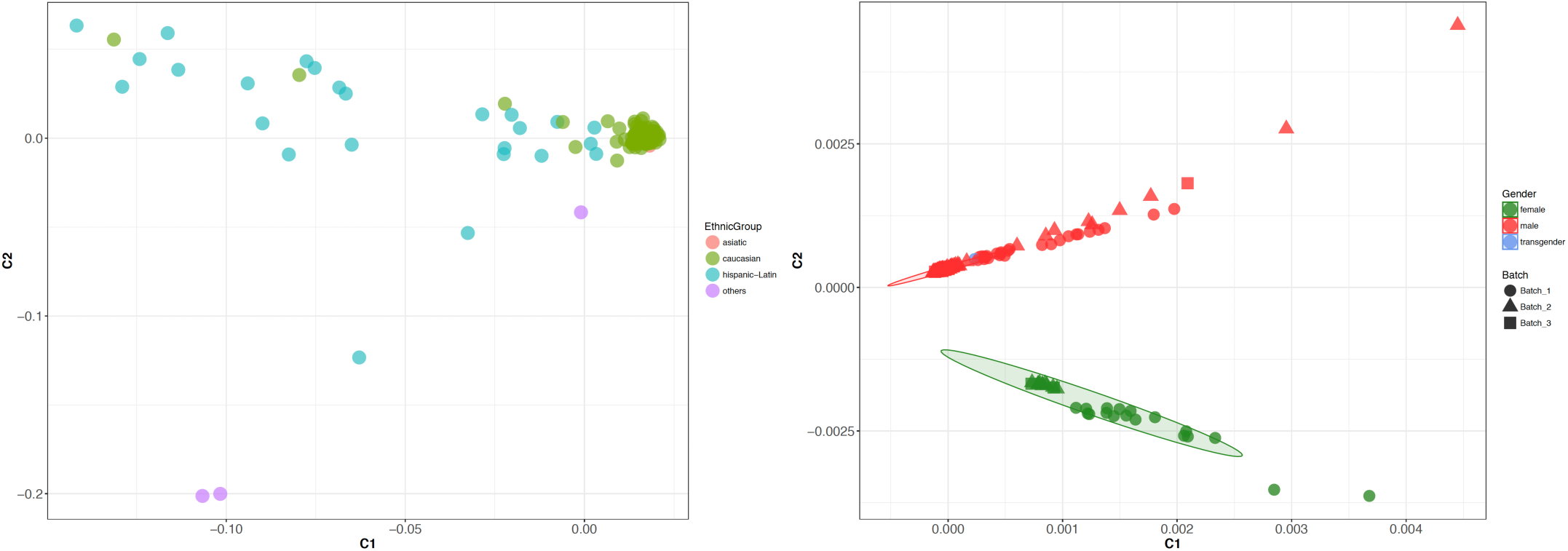
**a) Population stratification of the MetaHIV cohort based on IBD distances constructed from genetic variants matrix**. Individuals clustered according to their ancestry. Hispanic-latin subjects showed the maximum genetic variation among them because they descended from Spanish or portuguese-speaking cultures with mixed ancestry. **B) Multidimensional scaling (MDS) plot constructed from identity-by-missingness (IBM) distances**. Population stratification is caused by the effects of gender (f for females, m for males and t for transgender) on missingness, i.e. the proportion of missing genotypes is discordantly represented between females and males.

### Lack of significant associations between host genetics and gut microbiome composition

We assessed the interaction between QC-filtered genetic variants and three different gut microbial traits: gut enterotype (Bacteroides vs. Prevotella), methanogenic archaea (presence vs. absence) and gut microbial gene richness (as a continuous variable). Two independent logistic models were performed to test the association of genetic variants with the gut enterotype^23^ and the presence of methanogenic archaea^24^. A linear regression model was used for microbial gene richness. No significant associations were found between any of the three gut microbiome traits investigated and allelic variation under a canonical^31^ significance threshold of ~3×10^-7^.

### Exploratory analysis using less stringent criteria

With the purpose of providing a pilot exploration of host genomic variants more likely to be associated with the gut microbiome traits in our small dataset, we arbitrarily reduced the stringency of the p-value threshold to p <1e-5. Using this threshold, three genetic variants were linked to methanogenic detection and five were associated to microbial gene richness (Table 2, Figure 2, Supplementary Figure 2). None of them contributed to the enrichment of any reactome pathways.

**Table 2.**
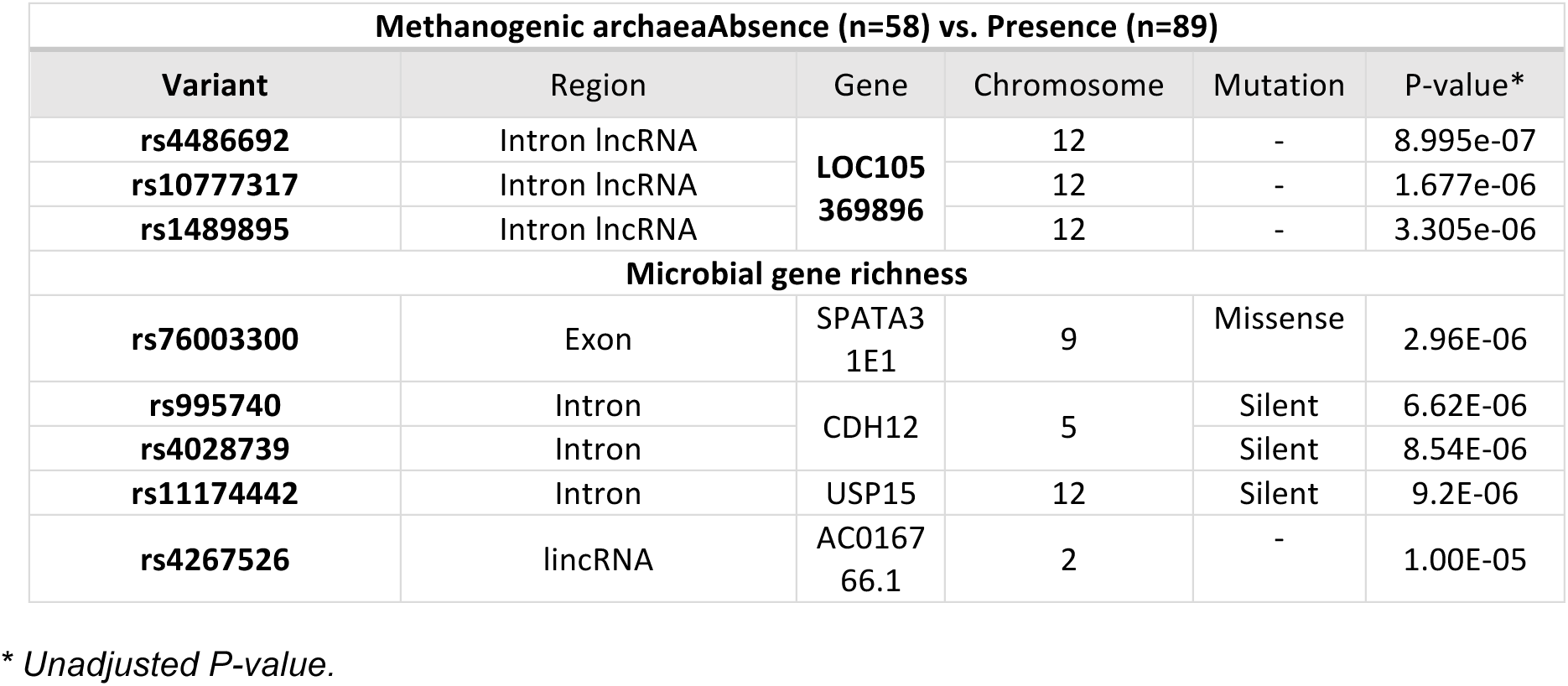
Genetic variants linked to methanogen detection and microbial gene richness using a p-value threshold <1e-5.

| Methanogenic archaeaAbsence (n=58) vs. Presence (n=89) |  |  |  |  |  |
| --- | --- | --- | --- | --- | --- |
| Variant | Region | Gene | Chromosome | Mutation | P-value* |
| rs4486692 | Intron lncRNA | LOC105369896 | 12 | - | 8.995e-07 |
| rs10777317 | Intron lncRNA |  | 12 | - | 1.677e-06 |
| rs1489895 | Intron lncRNA |  | 12 | - | 3.305e-06 |
| Microbial gene richness |  |  |  |  |  |
| rs76003300 | Exon | SPATA3 1E1 | 9 | Missense | 2.96E-06 |
| rs995740 | Intron | CDH12 | 5 | Silent | 6.62E-06 |
| rs4028739 | Intron |  |  | Silent | 8.54E-06 |
| rs11174442 | Intron | USP15 | 12 | Silent | 9.2E-06 |
| rs4267526 | lincRNA | AC016766.1 | 2 | - | 1.00E-05 |

\* *Unadjusted P-value.*

**Figure 2.**
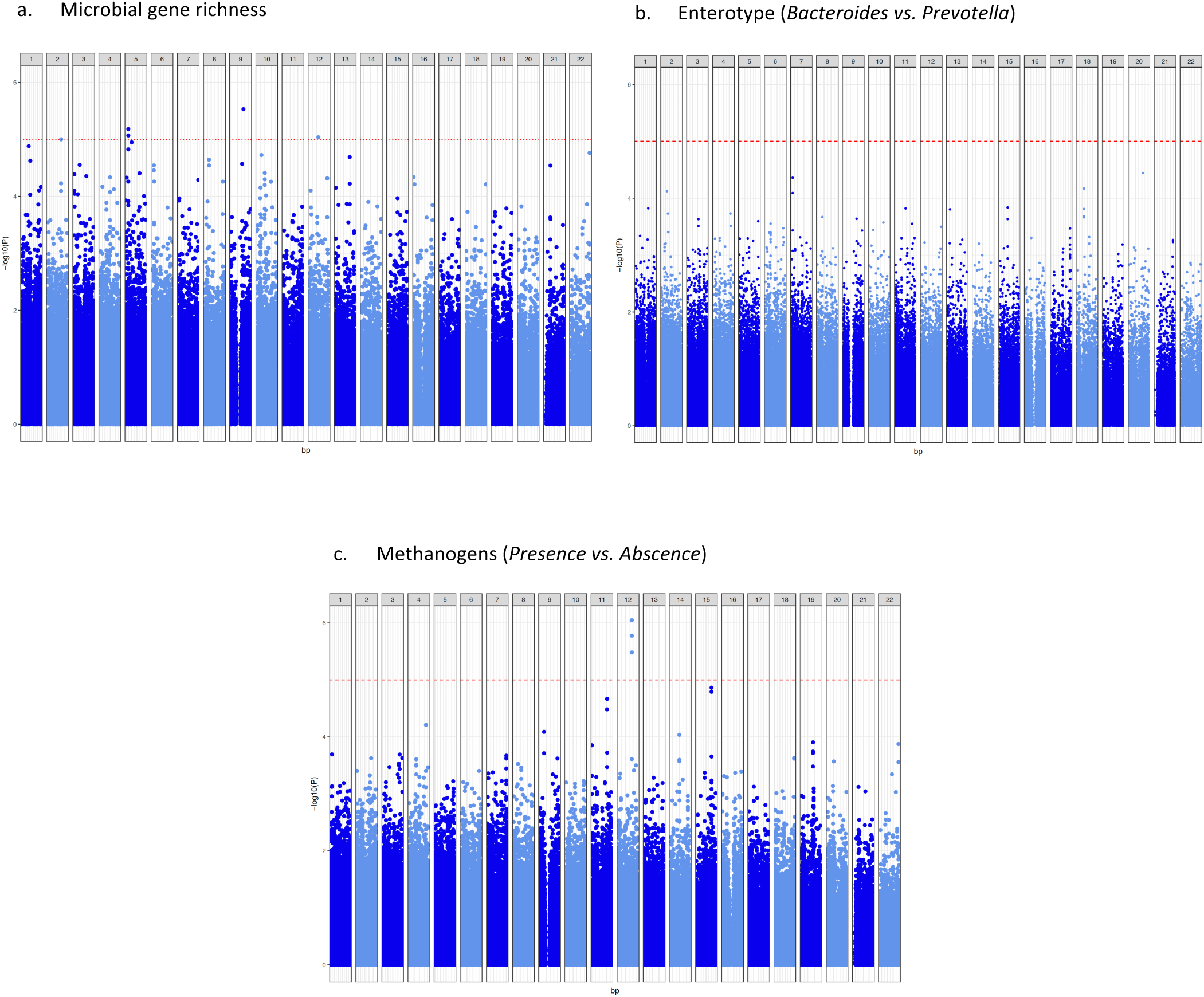
Association between host genetic variants and (a) gut microbial gene richness (b) enterotype and (c) detection of methanogens. Manhattan plots showing the correlation p-values (-log scale) for each filtered variant tested. Variants are represented by circles in their respective chromosome location (chromosome number indicated in the upper label). The significance threshold is indicated with a red dotted line. Variants depicted above this threshold are described in Table 2.

### Putative associations involving innate immunity and major histocompatibility genes

To evaluate the interaction between the gut microbiome composition and the innate immune system, we extracted from the InnateDB database^32^ the filtered host genetic variants from our cohort located in protein-coding sequences involved in human responses to microbes. Our dataset covered 6,127 variants extracted from InnateDB, which were distributed along 1,702 genes. None of these variants were associated to the explored microbiome features. To explore whether the MHC complex might be related with gut microbial traits, we investigated a total of 205 genetic variants distributed along 24 HLA genes. None of them were associated to screened microbial feature under a p-value threshold of p<1e-5.

## Discussion

Using a DNA-microarray-based genotyping strategy, we performed a pilot exploratory study to investigate whether the host genomic background could influence the gut microbiome composition in HIV-1-infected subjects ^23,24^. Following a GWAS-like analysis approach^31,33^, we found no statistically-significant associations between the covered host genetic background and Bacteroides vs. Prevotella clustering, detection of methanogens, or microbial gene richness. By setting a less stringent significant threshold, we identified a number of genetic associations with both methanogens presence and microbial gene richness. These should be seen as candidate associations that may exert small additive effects to the gut microbiome configuration and could be confirmed with properly-powered studies, including more participants and a genome-wide set of variants.

The small sample size of this study would only enable detecting variants associated with large effect sizes on gut microbiome composition. We thus suggest that, if the host genetics have any role in shaping the gut microbiome of people with or at risk of acquiring HIV-1, such effect is probably small. This is consistent with previous studies showing that the environment was, in fact, the main determinant of the gut microbiota composition in the general population^11^.

A study involving monozygotic and dizygotic human twins reported that the abundance of *Methanobrevibacter smithii* was more concordant between monozygotic than dizygotic twins, suggesting that the colonization of this group of microbes might have a genetic component^34^. In our study, the presence of methanogenic archaea was associated with three variants distributed along the same intron of LOC105369896, an uncharacterized long non-coding RNA (lncRNA) located in chromosome 12 (Table 2). Despite the vast majority of annotated lncRNA in the human genome have unknown functions, their participation in gene expression regulation, chromosome conformation, cell development and disease, has been widely reported^35^. A recent study in mice suggested that the lncRNA expression profile discriminates animals with different transplanted microbiota^36^. These results might be pinpointing to a putative role of lncRNA in human gut microbiota configuration, indicating that lncRNA could be involved in commensals recognition.

Microbial gene richness was the parameter with the higher number of linked genetic variants, although none of them clearly stood out. Two of these genetic variants are located within CDH12 gene, which encodes for cadherin-12 protein. Both of them are tagged as silent mutations, so we wouldn’t expect them to exert important phenotypic effects.

In mice, the major histocompatibility complex (MHC) was suggested to have an effect on the fecal microbiota composition by restricting the colonization of certain bacteria^37^. In our study, associations with HLA variants were not found. Nevertheless, in GWAS every association involving genes from the HLA complex should be interpreted with caution given the extreme polymorphism at both nucleotide and structural levels of this region^38^. Demonstration of the implication of HLA variation in human gut microbial configuration will thus require a specific well-powered study.

Unlike human cohorts, studies involving mice do allow for a better control of both genetic and environmental factors, that are important confounders^18^. In certain murine models, single mutations in genes implicated in innate immunity, including *leptin* or *toll-like receptor 5* (TLR5), have important effects on microbiome composition ^39,40^. While it is important to assess the contribution of host genotype in shaping the gut microbiota, it has been widely evidenced that the microbial responses induced by diet changes in mice, which are rapid and reproducible; dominate the host genotype effect ^41^.

This study has a number of limitations. First, its small sample size hinders our ability to detect statistically-significant associations once canonical false discovery corrections are implemented. Unlike in mouse models, testing genetic associations in humans require much larger and random sampling to compensate for environmental confounders such as diet and lifestyle habits. Our cohort included females and males from different geographic origins, which resulted in an expected population stratification^42^. However, the issue of population structure was controlled for using statistical methods developed to adjust for false associations^43,44^. Furthermore, data from the MetaHIV cohort should not be extrapolated to the general population because it is a very specific, not randomly-selected cohort of subjects who are either infected or at risk of being infected with HIV-1. Stool microbiome data may not necessarily reveal the effect of the genetic makeup on the dynamics of the microbial populations linked to intestinal mucosal surfaces^45^. Finally, no biological validation of any of the potential associations has been pursued^46^.

With all such caveats in mind, in this pilot exploratory analysis we do not find any consistent association between the host genetic makeup and the gut microbiota configuration in subjects who are either infected or at risk of becoming infected with HIV-1. Larger, adequately powered studies will be required to assess if host genetic variants mainly encoding for innate immunity genes, which shape the structure of epithelial cell surfaces ^47,18^, have an impact on the gut microbiota composition. In conclusion, prolonged immune deficiency and environmental and lifestyle factors play a much more important role than host genetics in shaping gut microbial communities in people living with HIV ^11,14,15,17,48^.

## Materials and Methods

### Study subjects

This study included subjects from MetaHIV, a cross-sectional study cohort of subjects infected or at risk of becoming infected with HIV-1 (see ^23^ for a detailed description of the cohort). HIV-1-infected subjects were recruited from two tertiary HIV-1 clinics in Barcelona, Catalonia, Spain. Most HIV-1-negative controls were enrolled from a prospective cohort of HIV-negative MSM who attend quarterly medical and counseling visits at a community-based center in Barcelona^49,50^.

### Ethics & Community Involvement

All subjects provided appropriate informed consent regarding genetic analyses. The study was reviewed and approved by the Institutional Review Boards of the Hospital Universitari Germans Trias i Pujol (reference PI-13-046) and the Hospital Vall d’Hebrón (reference PR(AG)109/2014). All participants provided written informed consent in accordance with the World Medical Association Declaration of Helsinki. The study concept, design, patient information and results were discussed with the HIVACAT Community Advisory Committee (CAC), who provided input on these aspects as well as on the presentation and dissemination of study results.

### Sample collection

Participants provided a single blood samples from which peripheral blood mononuclear cells (PBMCs) were isolated. DNA derived from PBMCs was extracted using the QIAamp^®^ DNA Blood minikit (QIAGEN Inc., Hilden, GER), and stored at -80ºC.

### Host genotyping

We genotyped 547,644 genetic variants from 147 subjects using the human Infinium^®^ CoreExome-24v1-0 BeadChip microarray and the Infinium^®^ HTS Assay Protocol Guide (Illumina Inc., San Diego, CA, USA). Exome genotyping was carried out at the *Institut de Medicina Predictiva del Cancer* (IMPPC). Genotype calling was done with the GenomeStudio Software version 2011.1 (Illumina Inc.). The average call rate was 99.9%, and no gender discrepancy was observed based on assay experiments. PLINK software v1.07^51^ was used to exclude individuals with >90% missing genotyping data (--mind 0.1) and variants not called in at least 10% of samples (--geno 0.1). Variants with minor allele frequency (MAF) lower than 0.01, and those out of Hardy-Weinberg equilibrium (P<1 x 10^-4^) were also removed (-hwe 0.0001). A total of 296,475 variants on 147 individuals were included in further analysis (Supplementary Figure 1). Variants passing such filters were annotated following the support data of the Infinium Core Exome kit provided by Illumina and the Ensembl Variant Effect Predictor tool^52^ (Table 1). Variants located in sexual chromosomes (chromosomes 23 and 24), pseudoautosomal regions (chromosome 25), mitochondrial DNA (chromosome 26) and control probes (chromosome 0) were excluded from the current analysis. Gene expression was assessed using the Human Protein Atlas database ^53^.

### Gut microbial characterization

Gut microbiota information was obtained from previously published studies using 16S rRNA^23^ and shotgun metagenomic sequencing^24^ of fecal samples from the same cohort. Based on those studies, we chose to investigate the association between the genetic makeup of the host and 3 specific gut microbiome traits, i.e.: (a) gut enterotype (Bacteroides vs. Prevotella), (b) detection of methanogenic archaea (*Methanobrevibacter smithii, M. unclassified* and *Methanosphaera stadtmanae*), and (c) gut microbial gene richness. Previous studies had suggested that HIV-1 infection was associated to shifts from Bacteroides to Prevotella predominance ^54–60^. Using 16S rRNA sequencing we previously observed, instead, that subjects enriched in Prevotella were mostly MSM, and there was no significant Bacteroides vs. Prevotella clustering by HIV-1 infection after controlling for HIV-1 risk group^23^. Using shotgun sequencing, we then found that low gene richness was linked to low nadir CD4+ T-cell counts and one of the most robust microbial markers of high microbial gene richness was, precisely, the detection of methanogenic archaea^24^.

### Statistical analysis

Population stratification was tested by constructing a matrix based on pairwise identity-by-state (IBS) distances. Identity-by-state distances were represented using a multidimensional scaling plot (Figure 1a). Effects of systematic batches and gender on missingness were assessed by clustering individuals according to their identity-by-missingness (IBM) distances, i.e. the proportion of missing genotypes discordantly missing by different participants (Figure 1b).

Genetic associations with both qualitative (gut enterotype, absence/presence methanogens) and quantitative microbial traits (microbial gene richness) were evaluated using logistic and linear regression models respectively, setting the additive version of the models and adding sex and gender as covariates. Statistic p-values were corrected using the Bonferroni adjustment to prevent false discovery, given the number of exonic variants tested and the cohort size. Associations between each microbial species abundance and genetic variation were not performed because the abundance of the microbes was not normally distributed, and substantial inflation of type I error rate was expected in such tests^61^.

A standard genome-wide significance threshold of ~3×10^-7^ for the European population has been previously proposed for simulated cohorts of N=100 with equal number of cases and controls ^31^. But, since 1) our population was stratified by ethnical origin; 2) the number of cases and controls was slightly unbalanced; and 3) we did not have a genome-wide coverage of genetic variants, variants with a minimum unadjusted p-value threshold of 1e-5 were considered for further analyses.

A gene set enrichment analysis of reactome pathways^62^ was performed for genes linked to significant variants using a binomial test and correcting p-values for multiple testing (Benjamini-hochberg procedure).

## Acknowledgements

This study was supported by philanthropic donations from Fundació Glòria Soler, the Gala SIDA 2015, 2016 editions, and People in Red – Barcelona 2017 edition. IrsiCaixa is supported by the RED de SIDA RD16/0025/0041 cofinanced by the ISCIII and the European Regional Development Fund (ERDF), “Investing in your future”. M.R. is funded through a FI-DGR grant (FI-B00184) from Agència de Gestió d’Ajuts Universitaris i de Recerca (AGAUR) at the Secretaria d’Universitats i Recerca del Departament d’Economia i Coneixement de la Generalitat de Catalunya. M.L.C. is funded through the grant MTM2015-64465-C2-1-R, Spanish Ministry of Economy and Competitiveness, Spain. We thank the members of the IGTP Genomics and Bioinformatics Core Facilities (Maria Pilar Armengol, Lauro Sumoy and Iñaki Martínez) for their contribution to this publication. The sponsors of the study had no role in study design, data collection, data analysis, data interpretation, or writing of the report. The corresponding author had full access to all study data, and had final responsibility for the decision to submit for publication.

## Author contributions

R.P., and B.C. conceived and designed the study. R.P., B.M., J.Co., J.S., J.N., C. B., M.Cr. and B.C recruited the study participants and performed their clinical evaluations. Fecal DNA was extracted, amplified and sequenced by M.P, M.Ca., and M.R. under the supervision of M.N. and R.P. Y.G, and J.R performed the bioinformatic and statistical analyses, with the supervision of M.N., C.B. and R.P.Y.G. and R.P wrote the paper, which was reviewed, edited and approved by all authors.

## Competing interests

The authors declare no competing interests.

## Data availability

The datasets generated during and/or analyzed during the current study are available in the National Center for Biotechnology Information -NCBI repository (Bioproject accession number: PRJNA307231, SRA accession number: SRP068240).

## Supplementary Material

**Supplementary Figure 1.** Workflow for the exploration of exonic variants linked to gut microbial composition in the MetaHIV cohort.

**Supplementary Figure 2.** Normal quantile-quantile plots (q-q plot) for P-values for a) gut microbial gene richness, b) enterotype and c) detection of methanogens.

## References

1. Cho, I. & Blaser, M. J. The human microbiome: at the interface of health and disease. Nat. Rev. Genet. 13, 260 (2012).

2. Human Microbiome Project Consortium. Structure, function and diversity of the healthy human microbiome. Nature 486, 207–14 (2012).

3. Wu, G. D. et al. Linking Long-Term Dietary Patterns with Gut Microbial Enterotypes. Science (80-. ). 334, 105–108 (2011).

4. Cotillard, A. et al. Dietary intervention impact on gut microbial gene richness. Nature 500, 585–588 (2013).

5. Morgan, X. C. et al. Dysfunction of the intestinal microbiome in inflammatory bowel disease and treatment. Genome Biol. 13, R79 (2012).

6. Serrano-Villar, S. et al. HIV infection results in metabolic alterations in the gut microbiota different from those induced by other diseases. Sci. Rep. 6, 26192 (2016).

7. Louis, P., Hold, G. L. & Flint, H. J. The gut microbiota, bacterial metabolites and colorectal cancer. Nat. Rev. Microbiol. 12, 661–672 (2014).

8. Forslund, K. et al. Disentangling type 2 diabetes and metformin treatment signatures in the human gut microbiota. Nature 528, 262 (2015).

9. Yatsunenko, T. et al. Human gut microbiome viewed across age and geography. Nature 486, 222 (2012).

10. Noguera-Julian, M. et al. Gut Microbiota Linked to Sexual Preference and HIV Infection. EBioMedicine 5, 135–146 (2016).

11. Rothschild, D. et al. Environment dominates over host genetics in shaping human gut microbiota. Nature 555, 210–215 (2018).

12. Ley, R., Hamady, M. & Lozupone, C. Evolution of mammals and their gut microbes. Science (80-.). 320, 1647–1651 (2008).

13. Goodrich, J. K., Davenport, E. R., Waters, J. L., Clark, A. G. & Ley, R. E. Cross-species comparisons of host genetic associations with the microbiome. Science 352, 532–535 (2016).

14. Zhernakova, A. et al. Population-based metagenomics analysis reveals markers for gut microbiome composition and diversity. Science 352, 565–569 (2016).

15. Gilbert, J. A. et al. Microbiome-wide association studies link dynamic microbial consortia to disease. Nature 535, 94–103 (2016).

16. Hall, A. B., Tolonen, A. C. & Xavier, R. J. Human genetic variation and the gut microbiome in disease. Nat. Rev. Genet. (2017). doi:10.1038/nrg.2017.63

17. Wang, J. et al. Genome-wide association analysis identifies variation in vitamin D receptor and other host factors influencing the gut microbiota. Nat. Genet. 48, 1396–1406 (2016).

18. Spor, A., Koren, O. & Ley, R. Unravelling the effects of the environment and host genotype on the gut microbiome. Nat. Rev. Microbiol. 9, 279–290 (2011).

19. Goodrich, J. K. et al. Genetic Determinants of the Gut Microbiome in UK Twins. Cell Host Microbe 19, 731–743 (2016).

20. Blekhman, R. et al. Host genetic variation impacts microbiome composition across human body sites. Genome Biol. 16, 191 (2015).

21. Goodrich, J. K. et al. Human genetics shape the gut microbiome. Cell 159, 789–799 (2014).

22. Hansen, E. E. et al. Pan-genome of the dominant human gut-associated archaeon, Methanobrevibacter smithii, studied in twins. Proc. Natl. Acad. Sci. U. S. A. 4599–606 (2011). doi:10.1073/pnas.1000071108

23. Noguera-Julian, M. et al. Gut Microbiota Linked to Sexual Preference and HIV Infection. EBioMedicine 5, 135–146 (2016).

24. Guillén, Y. et al. Low nadir CD4+ T-cell counts predict gut dysbiosis in HIV-1 infection. Mucosal Immunol. (2018).

25. Noguera-Julian, M. et al. Gut microbiota linked to sexual preference and HIV infection. EBioMedicine (in press), (2016).

26. Negredo, E. et al. Nadir CD4 T cell count as predictor and high CD4 T cell intrinsic apoptosis as final mechanism of poor CD4 T cell recovery in virologically suppressed HIV-infected patients: Clinical implications. Clin. Infect. Dis. 50, 1300–1308 (2010).

27. Bray, S. et al. Predictive value of CD4 cell count nadir on long-term mortality in HIV-positive patients in uganda. HIV/AIDS - Res. Palliat. Care 4, 135–140 (2012).

28. Arumugam, M. et al. Enterotypes of the human gut microbiome. Nature 473, 174–180 (2011).

29. Le Chatelier, E. et al. Richness of human gut microbiome correlates with metabolic markers. Nature 500, 541–546 (2013).

30. Sherry, S. T. et al. dbSNP: the NCBI database of genetic variation. Nucleic Acids Res. 29, 308–311 (2001).

31. Hoggart, C. J., Clark, T. G., Iorio, M. de, Whittaker, J. C. & Balding, D. J. Genome-Wide Significance for Dense SNP and Resequencing Data. Genet. Epidemiol. 32, 179–185 (2008).

32. Lynn, D. J. et al. InnateDB: Facilitating systems-level analyses of the mammalian innate immune response. Mol. Syst. Biol. 4, (2008).

33. Dudbridge, F. & Gusnanto, A. Estimation of significance thresholds for genomewide association scans. Genet. Epidemiol. 32, 227–234 (2008).

34. Turnbaugh, P. J. et al. A core gut microbiome in obese and lean twins. Nature 457, 480 (2008).

35. Quinn, J. J. & Chang, H. Y. Unique features of long non-coding RNA biogenesis and function. Nat. Publ. Gr. 17, 47–62 (2016).

36. Liang, L., Ai, L., Qian, J., Fang, J. & Xu, J. Long noncoding RNA expression profiles in gut tissues constitute molecular signatures that reflect the types of microbes. Nat. Publ. Gr. 1–8 (2015). doi:10.1038/srep11763

37. Toivanen, P., Vaahtovuo, J. & Eerola, E. Influence of Major Histocompatibility Complex on Bacterial Composition of Fecal Flora. Infect. Immun. 69, 2372–2377 (2001).

38. Kennedy, A. E., Ozbek, U. & Dorak, M. T. What has GWAS done for HLA and disease associations? Int. J. Immunogenet. 44, 195–211 (2017).

39. Ley, R. E. et al. Obesity alters gut microbial ecology. Proc. Natl. Acad. Sci. 102, 11070–11075 (2005).

40. Vijay-Kumar, M. et al. Metabolic Syndrome and Altered Gut Microbiota in Mice Lacking Toll-Like Receptor 5. Science 328, 228–231 (2010).

41. Carmody, R. N. et al. Diet dominates host genotype in shaping the murine gut microbiota. Cell Host Microbe 17, 72–84 (2015).

42. Tian, C., Gregersen, P. K. & Seldin, M. F. Accounting for ancestry: Population substructure and genome-wide association studies. Hum. Mol. Genet. 17, 143–150 (2008).

43. Clarke, G. M., Anderson, C. a, Pettersson, F. H., Cardon, L. R. & Andrew, P. Basic statistical analysis in genetic case-control studies. Nat. Protoc. 6, 121–133 (2011).

44. Bush, W. S. & Moore, J. H. Chapter 11: Genome-Wide Association Studies. PLoS Comput. Biol. 8, (2012).

45. Donaldson, G. P. et al. Gut microbiota utilize immunoglobulin A for mucosal colonization. Science (80-. ). eaaq0926 (2018). doi:10.1126/science.aaq0926

46. Williams, S. M. et al. Problems with genome-wide association studies. Science (80-. ). 5833, 1840–1842 (2007).

47. Dabrowska, K. & Witkiewicz, W. Correlations of host genetics and gut microbiome composition. Front. Microbiol. 7, 1–7 (2016).

48. Moschen, A. R., Wieser, V. & Tilg, H. Dietary factors: Major regulators of the Gut’s microbiota. Gut Liver 6, 411–416 (2012).

49. Coll, J. et al. Early diagnosis of HIV infectious and detection of asymptomatic STI in a community-based organization addressed to MSM. in 8th IAS Conference on HIV Pathogenesis Treatment and Prevention, 19-22 July 2015 (2015).

50. Meulbroek, M. et al. BCN Checkpoint, a community-based centre for men who have sex with men in Barcelona, Catalonia, Spain, shows high efficiency in HIV detection and linkage to care. HIV Med. 14, 25–28 (2013).

51. Purcell, S. et al. PLINK: A Tool Set for Whole-Genome Association and Population-Based Linkage Analyses. Am. J. Hum. Genet. 81, 559–575 (2007).

52. McLaren, W. et al. The Ensembl Variant Effect Predictor. Genome Biol. 17, 122 (2016).

53. Uhlén, M. et al. Tissue-based map of the human proteome. Science (80-. ). 347, (2015).

54. Monaco, C. L. et al. Altered Virome and Bacterial Microbiome in Human Immunodeficiency Virus-Associated Acquired Immunodeficiency Syndrome. Cell Host Microbe 19, 311–22 (2016).

55. Williams, B., Landay, A. & Presti, R. M. Microbiome alterations in HIV infection a review. Cell. Microbiol. 18, 645–51 (2016).

56. Williams, B. et al. A Summary of the First HIV Microbiome Workshop 2015. AIDS Res. Hum. Retroviruses (2016). doi:10.1089/AID.2016.0034

57. Ling, Z. et al. Alterations in the Fecal Microbiota of Patients with HIV-1 Infection: An Observational Study in A Chinese Population. Sci. Rep. 6, 30673 (2016).

58. Vázquez-Castellanos, J. F. et al. Altered metabolism of gut microbiota contributes to chronic immune activation in HIV-infected individuals. Mucosal Immunol. 8, 760–72 (2015).

59. Mutlu, E. A. et al. A compositional look at the human gastrointestinal microbiome and immune activation parameters in HIV infected subjects. PLoS Pathog. 10, e1003829 (2014).

60. Lozupone, C. A. et al. HIV-induced alteration in gut microbiota: driving factors, consequences, and effects of antiretroviral therapy. Gut Microbes 5, 562–70 (2014).

61. Schwantes-An, T.-H. et al. Type I error rates of rare single nucleotide variants are inflated in tests of association with non–normally distributed traits using simple linear regression methods. BMC Proc. 10, 62 (2016).

62. Croft, D. et al. Reactome: a database of reactions, pathways and biological processes. Nucleic Acids Res. 39, D691–D697 (2011).

